# Demographic Analysis of Mutations in Indian SARS-CoV-2 Isolates

**DOI:** 10.1101/2021.09.22.461342

**Authors:** Kushagra Agarwal, Nita Parekh

## Abstract

In this study we carried out the early distribution of clades and subclades state-wise based on shared mutations in Indian SARS-CoV-2 isolates collected (27^th^ Jan – 27^th^ May 2020). Phylogenetic analysis of these isolates indicates multiple independent sources of introduction of the virus in the country, while principal component analysis revealed some state-specific clusters. It is observed that clade 20A defining mutations C241T (ORF1ab: 5’ UTR), C3037T (ORF1ab: F924F), C14408T (ORF1ab: P4715L), and A23403G (S: D614G) are predominant in Indian isolates during this period. Higher number of coronavirus cases were observed in certain states, *viz*., Delhi, Tamil Nadu, and Telangana. Genetic analysis of isolates from these states revealed a cluster with shared mutations, C6312A (ORF1ab: T2016K), C13730T (ORF1ab: A4489V), C23929T, and C28311T (N: P13L). Analysis of region-specific shared mutations carried out to understand the large number of deaths in Gujarat and Maharashtra identified shared mutations defining subclade, I/GJ-20A (C18877T, C22444T, G25563T (ORF3a: H57Q), C26735T, C28854T (N: S194L), C2836T) in Gujarat and two sets of co-occurring mutations C313T, C5700A (ORF1ab: A1812D) and A29827T, G29830T in Maharashtra. From the genetic analysis of mutation spectra of Indian isolates, the insights gained in its transmission, geographic distribution, containment, and impact are discussed.

## Introduction

The coronavirus disease 2019 (COVID-19) pandemic is caused by the Severe Acute Respiratory Syndrome Coronavirus 2 (SARS-CoV-2), a betacoronavirus belonging to the Coronaviridae family. Large variation in the rate of infectivity and fatality due to COVID-19 is observed across different countries and a similar trend is observed across various states of India. To understand at the genetic level the role of acquired mutations in the circulating SARS-CoV-2 virus and their possible impact on the spread and virulence, a detailed analysis of Indian isolates obtained from GISAID (Shu and McCauley 2017) during the period 27^th^ Jan – 27^th^ May 2020 is carried out. During the early phase (Jan-Mar), positive cases were mainly due to individuals with travel history. The analysis of genetic variations accumulated is expected to indicate the impact of contact tracing, quarantine, and lockdown in containing the spread of COVID-19 during this period. A detailed state-wise distribution of shared mutations and their global distribution across the world is carried out to understand the transmission and virulence of the disease within and between states. This would help in identifying mutations responsible for large variation in the number of cases and deaths across different states in India and associated subclades of Indian isolates.

## Materials and Methods

For the period 27^th^ Jan – 27^th^ May 2020, 705 Indian SARS-CoV-2 viral isolates sequence data was obtained from GISAID (Shu and McCauley 2017). Of these 20 had incomplete metadata or low-quality genomes and were discarded and 685 isolates were considered for analysis. A total of 1279 variations were identified on comparing 685 Indian SARS-CoV-2 isolates with Wuhan-1 isolate as reference (summarized in **Supplementary File S1.xlsx**). Phylogenetic analysis of these isolates was carried out using Nextstrain resources Augur and Auspice (Hadfield *et al*. 2018). State-wise distribution of these variations was carried out to understand at the genetic level the high number of cases in the states of New Delhi, Tamil Nadu, Telangana, and high fatality rates in Gujarat and Maharashtra. To identify India-specific or state-specific variations, diversity (entropy) plots obtained using the Nextstrain resources were analysed at each site in the viral genome. State-specific clusters based on shared mutations during the initial period of the pandemic were identified by performing principal component analysis (PCA) on the mutational profile of Indian isolates. To assess the impact of India-specific non-synonymous mutations on protein stability, two machine learning based tools, I-Mutant2.0 (Capriotti *et al*. 2005) and MUpro (Cheng *et al*. 2006), were used. These tools predict both the direction of the change in protein stability and the associated DDG values (DDG = DG (wildtype) – DG (mutated protein)) in Kcal/mol, pH = 7, Temperature = 298K) for a protein structure or sequence. The DDG value > 0 indicates increased protein stability (I), while DDG < 0 corresponds to destabilizing effect of the mutation (D). In I-Mutant2.0, the sign of DDG is based on SVM classifier, and the associated confidence score is given by the reliability index. On the other hand, MUpro provides sign change prediction using two models, one SVM-based and the other using Neural Networks. In Table-1, the predicted sign of DDG by I-Mutant2.0 and MuPRO along with their respective confidence scores is reported. While we report individual effects of mutations on protein stability, some of the mutations in a haplotype may just be hitchhiking mutations.

## Results and Discussion

### Clade Analysis

Phylogenetic tree of Indian isolates constructed with respect to Wuhan-1 isolate as reference using the Nextstrain pipeline is shown in Figure 1. We observe that all the five clades (defined in the new Nextstrain classification scheme) were present in India during this early period with the following distribution: **19A**: 264 samples (38.54%), **19B**: 44 samples (6.42%), **20A**: 300 samples (43.80%), **20B**: 75 samples (10.95%), **20C**: 2 samples (0.29%). The earliest recorded entries of SARS-CoV-2 in the country are both from Kerala with a travel history from Wuhan, China. Acc. ID: EPI_ISL_413522 on 27^th^ Jan 2020 belonging to the root clade 19A and a sample of clade 19B on 31^st^ Jan 2020 (Acc. ID: EPI_ISL_413523). By January end, these two clades had globally spread across most parts of the world, including India. The earliest sample corresponding to clade 20A is dated 3^rd^ March, of a tourist from Italy (Acc. ID: EPI_ISL_420543). Two samples corresponding to clade 20B were observed during the same time, one having contact with another Indian with travel history from Italy (Acc. ID: EPI_ISL_426179), dated 2^nd^ March, and the other on 29^th^ February (Acc. ID: EPI_ISL_414515), with no state or travel history available. Two samples of clade 20C were observed (Acc. ID: EPI_ISL_435051 and EPI_ISL_435052), dated 13^th^ April in Gujarat with no travel history or contact with anyone with travel history. These results clearly indicate early community spread of the virus in the country.

**Fig. 1.**
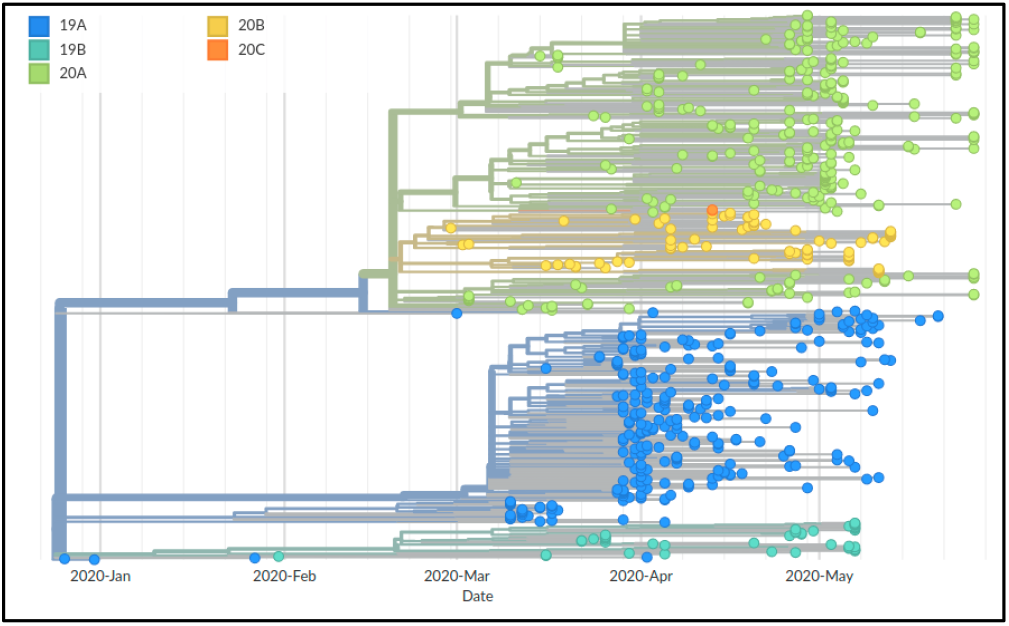
Phylogenetic tree obtained with Wuhan-1 isolate as reference depicts the divergence of Indian SARS-COV-2 isolates during the period 27^th^ Jan – 27^th^ May 2020.

Highest number of isolates belong to clade 20A (43.80%) in accordance with the global trend, with the next large cluster corresponding to root clade 19A (38.54%), comprising ∼82% of reported isolates in India till 27^th^ May. Our analysis revealed subclusters of clade 20A and 19A, respectively, with defining mutations. It would be interesting to study the state-wise distribution of this clade along with travel history to assess any community transmission during the lockdown.

### State-wise Distribution of Clades

Of 685 Indian isolates, 658 isolates were considered for state-wise distribution analysis (**Supplementary Figure 1**) as 27 isolates had no state information available. Clade 20A is predominantly observed in Gujarat (178/201), followed by West Bengal (30/45), Odisha (22/46), Madhya Pradesh (16/19), and Maharashtra (16/80), with the earliest reported sample from Gujarat dated 5^th^ April (Acc. ID: EPI_ISL_426414). Multiple independent entries of this clade are indicated based on clusters with shared mutations (discussed later). The second major clade 19A is predominant in Telangana (75/97), followed by Delhi (55/76), Maharashtra (31/80), and Tamil Nadu (19/34). Very few isolates of clade 19B (Odisha (17), Maharashtra (10), Gujarat (8) and West Bengal (5)), clade 20B (Maharashtra (23), Telangana (15), Tamil Nadu (15), and Delhi (7)) and clade 20C (only from Gujarat (2)) are observed. We observed that the genomic sequences of a few isolates from Odisha were shorter in length with missing bases; 1 - 29 bases in 5’ UTR (38 isolates) and 29686 - 29903 bases in 3’ UTR (39 isolates), which are likely to be due to sequencing artefact as majority of these are from the same sequencing laboratory. The isolates from Odisha also contain other deletion regions, 23842 - 24400 bases in S gene, 26306-26524 bases in E gene, 27527 – 28033 bases covering ORF7b and ORF8 genes in 8 samples, and 28462 - 28680 bases in N gene (10 samples), 29000 – 29685 bases covering N and ORF10 genes (15 samples), which may be region-specific.

### Most frequent mutations in Indian isolates

The important mutations in Indian isolates and their distribution state-wise is summarized in Table 1. Mutation C241T in the 5’ UTR region of ORF1ab is the most common mutation with incidence in more than half of the Indian isolates. It is highly prevalent in Gujarat (183/201), followed by Maharashtra (39/80) and West Bengal (33/45). Being a non-genic mutation, it does not cause an amino acid change. However, being part of the stem-loop region upstream of the ORF1ab gene, it may involve in differential RNA binding affinity to the ribosome and affect translational efficiency of the virus, though no change in the RNA structure was observed for the mutant form (Alam *et al*. 2021). Chaudhari *et al* (2021) analysed the role of C241T mutation in the interaction of stem-loop region with the host replication factors, MADP1 Zinc finger CCHC-type and RNA-binding motif 1 (hnRNP1). Molecular docking and molecular dynamics simulations suggest reduced replication efficiency of the virus within host and may result in less mortality.

**Table 1.** Important mutations observed in Indian samples along with their frequency, state distribution, and I-Mutant2.0 and MUpro SVM scores. Acronyms used for states: GJ - Gujarat, WB – West Bengal, MH – Maharashtra, TL – Telangana, DL – Delhi, TN – Tamil Nadu.

| Variant Description | Nucleotide Mutation | Protein | Amino Acid Mutation | Frequency |  | I-Mutant2.0 Sign (Reliability Index) | MUpro Sign (Confidence score) |  |
| --- | --- | --- | --- | --- | --- | --- | --- | --- |
|  |  |  |  | India | States |  | SVM | NN |
| Clade 20A | C241T | 5' UTR | - | 378 (55.2) | GJ (91.0), WB (73.3), MH (48.8) | - | - | - |
| Maharashtra-specific | C313T | ORF1ab (nsp1) | - | 39 (5.7) | MH (25.0) | - | - | - |
| Subclade I/GJ-20A | C2836T | ORF1ab (nsp3) | - | 51 (7.4) | GJ (25.4) | - | - | - |
| Clade 20A | C3037T | ORF1ab (nsp3) | - | 374 (54.6) | GJ (90.6), WB (73.3), MH (48.8) | - | - | - |
| Maharashtra-specific | C5700A | ORF1ab (nsp3) | A1812D | 33 (4.8) | MH (26.2) | D (4) | I (0.52) | D (-0.67) |
| Co-occurs with subclade I/A3i | C6310A | ORF1ab (nsp3) | S2015R | 44 (6.4) | TL (15.5), DL (10.5) | D (3) | I (0.50) | I (0.52) |
| Subclade I/A3i | C6312A | ORF1ab (nsp3) | T2016K | 228 (33.3) | TL (73.2), DL (69.7), TN (55.9) | D (7) | D (-1) | D (-0.78) |
| Co-occurs with subclade I/A3i | G11083T | ORF1ab (nsp6) | L3606F | 258 (37.7) | TL (72.2), DL (71.0), TN (55.9) | D (7) | D (-1) | D (-0.97) |
| Subclade I/A3i | C13730T | ORF1ab (nsp12) | A4489V | 239 (34.9) | TL (74.2), DL (71.0), TN (55.9) | I (1) | I (0.36) | D (-0.52) |
| Clade 20A | C14408T | ORF1ab (nsp12) | P4715L | 367 (53.6) | GJ (88.6), WB (73.3), MH (46.2) | D (6) | I (0.34) | I (0.78) |
| Subclade I/GJ-20A | C18877T | ORF1ab (nsp14) | - | 126 (18.4) | GJ (52.2) | - | - | - |
| Co-occurs with subclade I/A3i | C19524T | ORF1ab (nsp14) | - | 60 (8.8) | TL (20.6), DL (13.2) | - | - | - |
| Subclade I/GJ-20A | C22444T | S | - | 71 (10.4) | GJ (31.8) | - | - | - |
| Clade 20A | A23403G | S | D614G | 373 (54.4) | GJ (89.6), WB (73.3), MH (48.8) | D (7) | D (-0.57) | D (-0.98) |
| Subclade I/A3i | C23929T | S | - | 224 (32.7) | TL (71.1), DL (69.7), TN (47.0) | - | - | - |
| Subclade I/GJ-20A | G25563T | ORF3a | Q57H | 134 (19.6) | GJ (53.7) | D (7) | D (-0.02) | D (-0.56) |
| Subclade I/GJ-20A | C26735T | M | - | 123 (18.0) | GJ (50.2) | - | - | - |
| Subclade I/A3i | C28311T | N | P13L | 236 (34.4) | TL (76.0), DL (68.4), TN (55.9) | D (4) | D (-0.08) | D (-0.82) |
| Subclade I/GJ-20A | C28854T | N | S194L | 73 (10.6) | GJ (32.8) | I (3) | I (0.41) | I (0.88) |
| Clade 20B | G28881A | N | R203K | 75 (10.9) | TN (44.1), MH (28.8), TL (15.5) | D (5) | D (-0.59) | D (-0.88) |
| Clade 20B | G28882A | N | R203K | 75 (10.9) | TN (44.1), MH (28.8), TL (15.5) | D (5) | D (-0.59) | D (-0.88) |
| Clade 20B | G28883C | N | G204R | 75 (10.9) | TN (44.1), MH (28.8), TL (15.5) | D (7) | I (0.50) | I (0.76) |
| Maharashtra-specific | A29827T | 3' UTR | - | 44 (6.4) | MH (55.0) | - | - | - |
| Maharashtra-specific | G29830T | 3' UTR | - | 61 (8.9) | MH (73.8) | - | - | - |

The D614G mutation (A23403G) in the Receptor Binding Domain (RBD) of Spike protein has been widely studied globally and has a similar distribution in the Indian samples. It was first observed in China and Germany in late January and is now the most prevalent mutation worldwide. It is observed with over 50% representation (373/685) in Indian isolates, and with a high predominance in Gujarat (180/201). Glycine being less bulky than aspartic acid, contributes to a more flexible hinge region in the Spike protein, enabling more efficient receptor-ligand binding. This provided selective advantage to the virus in infection and transmission, making it predominant all over the world (Zhang *et al*. 2020). Plante *et al*. (2020) characterized the replication of D614 and G614 viruses in a primary human airway tissue culture model and showed that the mutation enhances viral replication through increased virion infectivity and enhances the stability of the virus. Another study (Daniloski *et al*. 2021) suggested possible mechanism for increased infectivity of G614 variant to it being more resistant to proteolytic cleavage. This stability confers G614 variant efficient transmission ability. Protein stability analysis using I-Mutant2.0 and MUpro indicate that the mutation has a destabilizing effect due to increased flexibility provided by glycine. The D614G mutation is found to co-occur with C241T, C3037T, and C14408T (Korber *et al*. 2020) and these are the defining mutations for clade 20A in Nextstrain and are predominantly found in North America and Europe. The C14408T mutation within the RNA-dependent RNA polymerase (RdRp) encoding region of ORF1ab is a missense mutation that leads to an amino acid change from Proline to Leucine (P4715L). It is found in Nsp12 region which is involved in replication and pathogenesis and is a potential target for antiviral candidates in coronaviruses. It is also observed in >50% of Indian samples with predominance in Gujarat (178/201), Maharashtra (37/80), and West Bengal (33/45). From Table 1 it may be noted that I-Mutant2.0 analysis predicts decreased stability for the C14408T mutation while MUpro scores indicate stabilizing effect. Nucleotide mutation C3037T is a silent mutation in the Nsp3 protein of ORF1ab (F924F) and no functional significance of this mutation is reported. The haplotype defined by these four co-occurring mutations is now globally the dominant form. This haplotype was proposed to be related to the pathogenicity of the virus and correlated with high death rates in Europe (Bai *et al*. 2020). High prevalence of these mutations in Gujarat with similar death rates suggests probable transmission of the virus to Gujarat from North America and Europe. Its containment in the state due to lockdown.

Clade 20B defining mutations, G28881A, G28882A, and G28883C, results in two adjacent amino acid changes R203K and G204R in the nucleocapsid (N) protein. This trinucleotide-bloc mutation, 28881-28883: GGG > AAC, is reported to result in two sub-strains of SARS-CoV-2, *viz*., SARS-CoV-2g and SARS-CoV-2a. The AAC genotype is observed from March globally and in India (75 samples) it is mainly observed in Maharashtra (23/80), Tamil Nadu (15/34), and Telangana (15/97). Mutation RG > KR is observed to disrupt the S-R motif by introducing Lysine between them. This may affect phosphorylation of the SR-rich domain that plays an important role in the cellular localization and translation inhibitory function of the N protein (Ayub 2020). Earlier experimental work on SARS-CoV-1 has shown reduced pathogenicity on deleting part of the SR domain (Tylor *et al*. 2009). These mutations also reduce the number of miRNA binding sites from seven to three and thereby increase the likelihood of viral infection (Maitra *et al*. 2020). Stability analysis using both I-Mutant2.0 and MUpro indicate decrease in protein stability for R203K mutation. However, for G204R mutation, I-Mutant2.0 predicts a decrease in stability while MUpro (both models) indicate increase in stability, which is understandable as Gly has no side chains. Since these are adjacent mutations, it would make sense to analyse the impact of the dipeptide RG > KR mutation. High prevalence of these mutations in Gujarat with similar death rates suggest probable transmission of the virus to Gujarat from North America and Europe. Its containment in the state due to lockdown and local transmission is probably the reason for its prevalence in Gujarat state.

### Mutations specific to Indian isolates

The diversity plots for SARS-CoV-2 isolates from India and globally are given in Figure 2 (a) and 2 (b), respectively. Sites exhibiting higher entropy in Indian isolates compared to global isolates are marked in Figure 2 (a) and results are also summarized in Table 1. Four of these, C6312A (T2016K) and C13730T (A4489V) in ORF1ab, C23929T in Spike protein, and C28311T (P13L) in N gene have been reported in an earlier study as India specific subclade I/A3i defining mutations (Banu *et al*. 2020). Two other India-specific mutations, C6310A (S2015R) and C19524T in ORF1ab are identified to be associated with this subclade and correspond to two branch-points of subclade I/A3i with predominance in Delhi and Telangana.

**Fig. 2.**
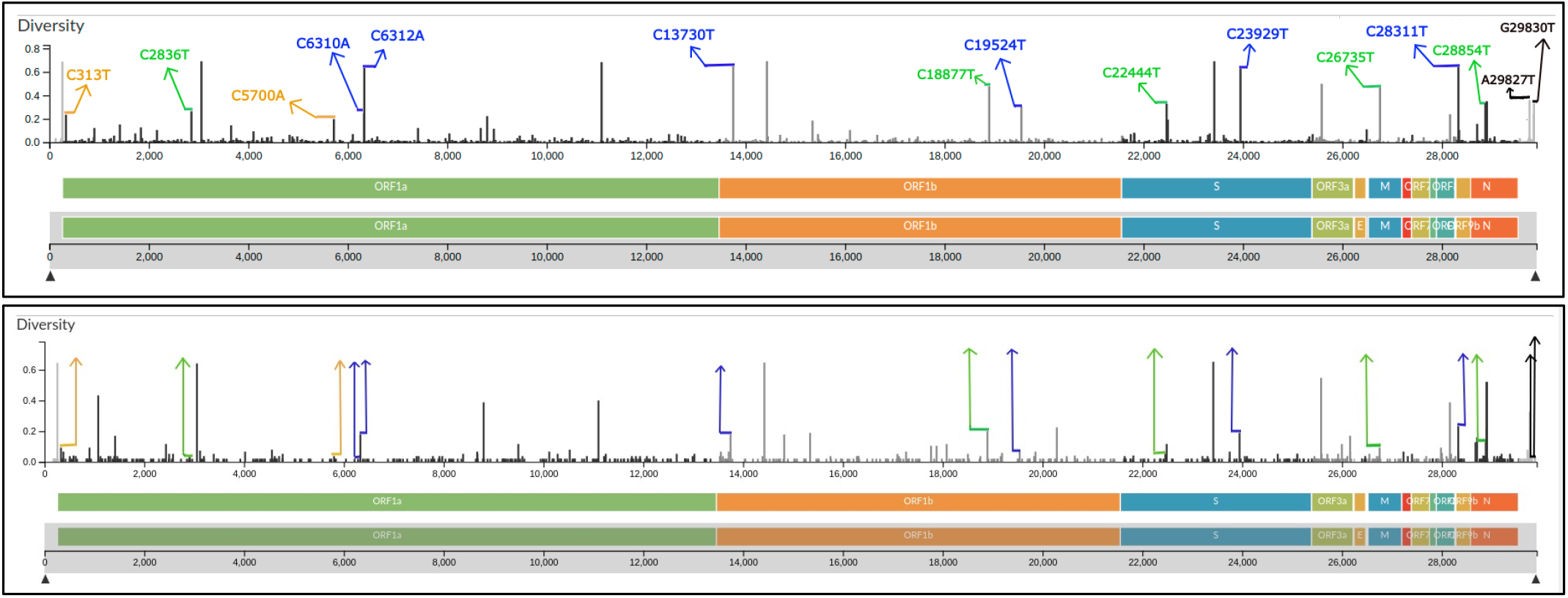
The diversity plots shown for isolates from (a) India and (b) World. In (a) 15 mutations predominant in India are marked in: Blue (clade 19A): C6310A, C6312A, C13730T, C19524T, C23939T and C28311T, Green (clade 20A): C2836T, C18877T, C22444T, C26735T, C28854T, Orange (clade 20B): C313T, C5700A, and Black (not specific to any particular clade): A29827T, G29830T.

Our analysis also revealed mutations C313T (synonymous) and C5700A (A1812D) in ORF1ab which co-occur with high frequency in India compared to their global presence. These mutations define a unique cluster of viral isolates predominantly observed in Maharashtra and Telangana states. The mutation C5700A is observed to branch out of clade 20B primarily, and some clade 20A samples also exhibit this mutation (**Supplementary Figure 2a**). This mutation is unique to the Indian region (∼5% frequency) with no presence globally (Figure 2(a) and 2(b)) and its co-occurrence with mutation C313T is observed in 33 Indian samples: Maharashtra (21), Telangana (8) and Gujarat (4). The first instance of this mutation was on 4^th^ April 2020 in Maharashtra, and it is also reported in an earlier study on 90 sequences from western India (Paul *et al*. 2020). Its presence is observed to increase with 10% of Indian samples by July 2020 carrying this mutation, indicating local transmission. Mutation C5700A lies in the Nsp3 region of ORF1ab, which forms multi-subunit assemblies with other non-structural proteins for the formation of replication-transcription complex (Lei *et al*. 2018). It has the ability to alter surface electrostatic environment in the proximity of Nsp3’s viral protease domain (Gupta *et al*. 2021). It is shown to exhibit a destabilizing effect on the protein by I-Mutant2.0 and MUpro NN model (Table 1). In Figure 2(a), the mutations in green, C2836T, C18877T, C22444T, C26735T, C28854T are specific to Gujarat viral isolates while those in black, A29827T and G29830T are observed in Maharashtra, and their significance is discussed below.

### Geographical distribution of Indian subclade I/A3i

About one-third of Indian isolates (219/685) defined by co-occurring mutations C6312A, C13730T, C23929T, and C28311T, are associated with subclade I/A3i that branches out of clade 19A (**Supplementary Figure 2b**). State-wise distribution analysis of this subclade (**Supplementary Figure 3a**) revealed its predominance in Telangana (69/97, ∼71%), Delhi (52/76, ∼68%), Haryana (11/11) and Tamil Nadu (16/34). The mutation C13730T (ORF1ab: A4489V) lies in the RdRp protein’s NiRAN domain that is involved in RNA binding and nucleotidylation and is essential for viral replication (Gao *et al*. 2020). The nucleocapsid protein (N) is required for viral RNA replication, transcription, as well as genome packing (Hsin *et al*. 2018; Masters 2019). Mutation C28311T (N: P13L) is located in the protein’s intrinsically disordered region and may impair the terminal domain’s RNA-binding function (Chang *et al*. 2009; Chang *et al*. 2014). A previous study evaluated how this mutation may alter protein-protein interaction and proposed its impact on virus stability, potentially contributing to lower pathogenesis (Oulas *et al*. 2021). From the stability analysis, the two non-synonymous mutations representing subclade I/A3i (C6312A and C28311T) decrease the stability. Mutation C13730T is likely to decrease protein stability as indicated by the confidence score of MUpro (NN). For C6310A, the mutation is expected to increase the stability according to MUpro models, as the reliability score of I-Mutant is low.

Analysis of subclade I/A3i is important both scientifically and epidemiologically as its defining mutations are found in ∼32% of Indian samples, while outside India its distribution is very low (∼3.5%). This clearly hints at early community transmission due to some super spreader event during March-April, as it is highly unlikely that around one-third of the samples sharing the same set of mutations could have arisen by multiple independent entries with international travel history, especially when its presence globally is negligibly small. The first reported entry of this clade in India is on 16^th^ March 2020 in Telangana from an Indonesian citizen visiting India and globally first incidence of this subclade is in a sample from Saudi Arabia, dated 7^th^ March 2020. It is predominantly found in Asia (mostly Singapore and Malaysia) (**Supplementary Figure 3b**). The coincidence of subclade I/A3i in the country following Tablighi congregation held during early March, first reported case from Saudi Arabia on 7^th^ March, and several Indonesian citizens identified with these mutations in Delhi during that period, all hint at the Tablighi congregation event as the likely cause of spread of this subclade. No isolates belonging to this subclade were observed after May, except for 5 samples in India with the last one dated 13^th^ June 2020 (according to data available in Nextstrain). This indicates that spread of subclade I/A3i had been largely contained during lockdown with efforts of contact tracing and quarantine of infected individuals. Similarly, no further global spread of this subclade is observed, probably due to air travel ban, the efficacy of contact tracing and quarantining of COVID-19 positive individuals. Thus, genetic analysis can help in identifying the chain of transmission of infection and the success of measures used in its containment.

### Analysis of samples with travel history from foreign nations

With the ban on international flights from March 23^rd^, 2020 and lockdown imposed in the country two days later, no new transmissions were likely from outside the country. Further, lockdown would have promoted accumulation of region-specific mutations. Analysis of shared mutation sets can help in understanding the extent of local transmission and containment of the circulating strains in a localised region. In 14 samples with travel history from Iran, 13 samples had the mutations, G1397A, T28688C, G29742T. Amongst them, 11 also had the mutations, C884T, G8653T, A29879G, A29883T, A29901T. These mutations were observed in very few other Indian isolates; Ladakh (6/6), Kargil (1/1), and Maharashtra (1/80) indicating the efficacy in quarantining and contact tracing during the lockdown period. The 11 samples with travel history from Italy contained the mutations C3037T, C241T, C14408T (ORF1ab: P4175L), and A23403G (S: D614G), which are clade 20A defining mutations (in Nextstrain). These mutations are found in 50% of Indian samples and with highest frequency in Gujarat (∼90% samples). Eight samples from Indonesia sampled in Delhi around the time the Tablighi Jamaat religious congregation was conducted, contained the four defining mutations of subclade I/A3i, C23929T, C6312A, C13730T, and C28311T. As discussed above, this mutation set was mainly observed in Telangana, Delhi, and Tamil Nadu but no further increase was seen in the number of samples containing it.

### PCA Analysis of Indian SARS-CoV-2 isolates

Since travel between states was restricted because of lockdown, increase in number of samples with shared mutations is expected because of local community transmission. To see if any state-specific clustering is observed during this initial period, principal component analysis (PCA) on the mutational profile of Indian isolates was performed. Figure 3 shows the plot of the first two principal components that captured over 35% variance in this high-dimensional data. Mutations were sorted based on their loading scores to assess their impact on PC1. Top 10 mutations thus identified are: A23403G (Spike: D614G), C3037T, C241T, C14408T (ORF1ab: P4715L), G11083T (ORF1ab: L3606F), C28311T (N: P13L), C6312A (ORF1ab: T2016K), C13730T (ORF1ab: A4489V), C23929T, and G25563T (ORF3a: Q57H). Not surprisingly, these are also the top 10 most common mutations in Indian isolates. It may be noted in Figure 3 that the Gujarat samples shown in ‘pink’ form a distinct cluster. Samples from Telangana, Delhi, and Haryana states cluster to the right in the plot due to the shared mutation profile of subclade I/A3i. Samples from Maharashtra are scattered throughout the plot, though in closely grouped clusters. A detailed analysis revealed two sets of co-occurring mutations in Maharashtra isolates (discussed later).

**Fig. 3.**
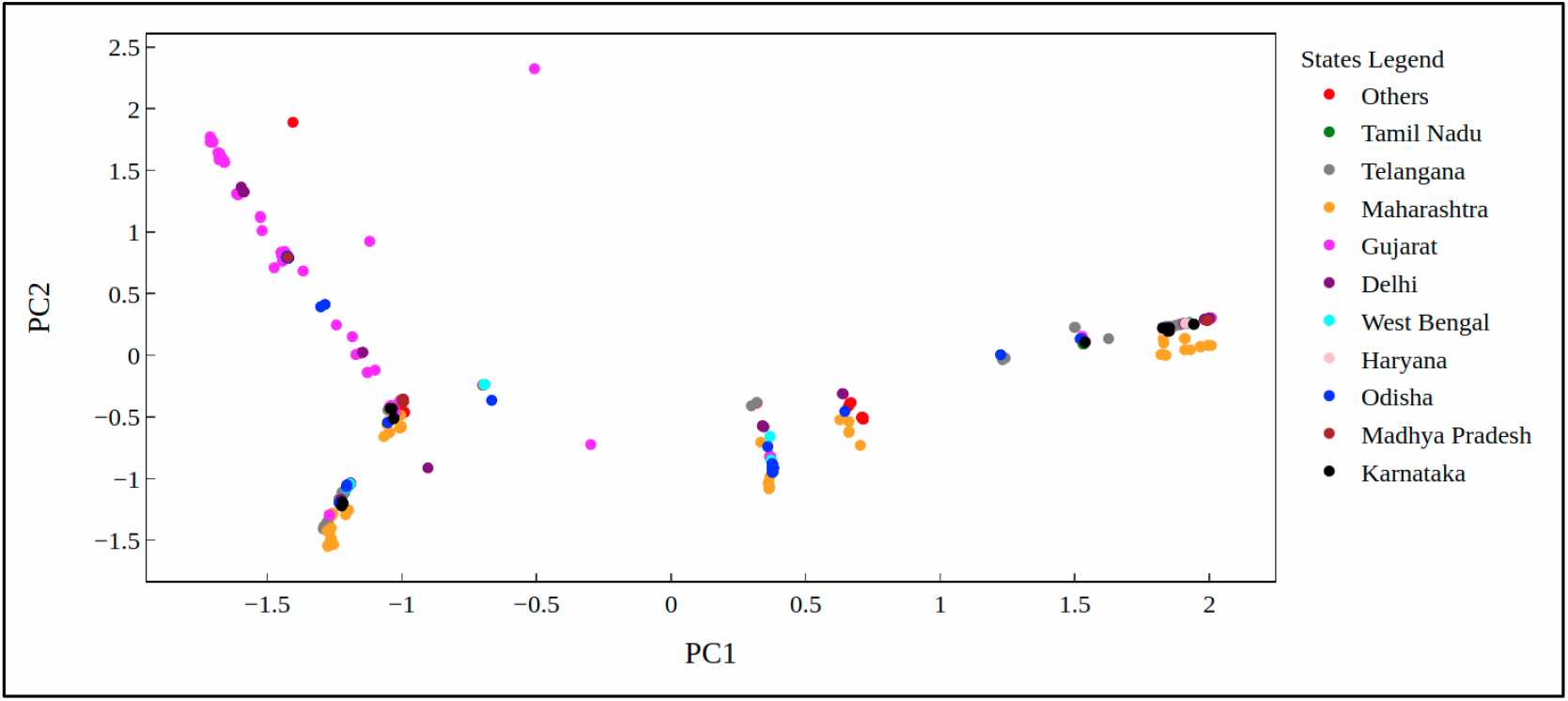
PCA plot of 685 samples coloured state-wise.

### Mutational Profile of Gujarat vs Rest of India

It is observed that certain countries like Italy, UK, Spain, etc. had large number of mortality cases. Severity of the circulating strains and large aging population were proposed to be the probable cause. India being a very vast country, it would be interesting to study if a similar pattern is observed across its different states. We analysed the infection and death rates across states and states with high severity of COVID-19 cases are discussed below. We observe Gujarat (5.12%) and Maharashtra (4.19%) reported higher death rates compared to country average (2.67%), as on 11^th^ July 2020. Recorded number of deaths in Gujarat is 2008 out of a total of 39194 cases while Karnataka (31105), Telangana (30946), and Uttar Pradesh (32363) with similar number of cases, recorded much fewer fatalities, 486, 331, and 862, respectively. Even some of the worst-hit regions such as Delhi (3.29%) and Tamil Nadu (1.39%) had much lower death rates compared to Gujarat. To understand this significantly large difference in the percentage of deaths in Gujarat at the genetic level, a detailed analysis of the mutational profile of sequences from Gujarat with that of Rest of India (RoI) was carried out. Table 2 summarizes non-synonymous mutations that are over- and under-represented in Gujarat isolates compared to the rest of country. Clade 20A defining mutations are observed in ∼90% of Gujarat samples. Due to countrywide lockdown from 25^th^ March 2020 clade 20A and its sub-clusters were localized in the state and are identified as Gujarat-specific mutations in Table 2. A reverse scenario, that is, under-representation of certain mutations is observed in Gujarat that has high frequency in RoI, e.g., mutations defining subclade I/A3i and clade 20B.

**Table 2.** Over- and under-represented mutations observed in SARS-CoV-2 isolates from Gujarat (201 samples) compared to the Rest of India, RoI (484 samples) are listed.

| Nucleotide Mutation | Protein | Amino Acid mutation | Count in Gujarat | Frequency in Gujarat (%) | Count in RoI | Frequency in RoI (%) |
| --- | --- | --- | --- | --- | --- | --- |
| *Mutations over-represented in Gujarat* |  |  |  |  |  |  |
| C241T | 5' UTR | - | 183 | 91.0 | 195 | 40.3 |
| C3037T | ORF1ab (nsp3) | - | 182 | 90.6 | 192 | 39.7 |
| A23403G | S | D614G | 180 | 89.6 | 193 | 39.9 |
| C14408T | ORF1ab (nsp12) | P4175L | 178 | 88.6 | 189 | 39.0 |
| G25563T | ORF3a | Q57H | 108 | 53.7 | 26 | 5.4 |
| C18877T | ORF1ab (nsp14) | - | 105 | 52.2 | 21 | 4.3 |
| C26735T | M | - | 101 | 50.2 | 22 | 4.5 |
| C28854T | N | S194L | 66 | 32.8 | 7 | 1.4 |
| C22444T | S | - | 64 | 31.8 | 7 | 1.4 |
| C2836T | ORF1ab (nsp3) | - | 51 | 25.4 | 0 | 0.0 |
| *Mutations under-represented in Gujarat* |  |  |  |  |  |  |
| C313T | ORF1ab (nsp1) | - | 2 | 1.0 | 37 | 7.6 |
| C5700A | ORF1ab (nsp3) | A1812D | 4 | 2.0 | 29 | 6.0 |
| C6310A | ORF1ab (nsp3) | S2015R | 6 | 3.0 | 38 | 7.8 |
| C6312A | ORF1ab (nsp3) | T2016K | 12 | 6.0 | 216 | 44.6 |
| G11083T | ORF1ab (nsp6) | L3606F | 16 | 8.0 | 242 | 50.0 |
| C13730T | ORF1ab (nsp12) | A4489V | 12 | 6.0 | 227 | 46.9 |
| C19524T | ORF1ab (nsp14) | - | 6 | 3.0 | 54 | 11.2 |
| C23929T | S | - | 12 | 6.0 | 212 | 43.8 |
| C28311T | N | P13L | 13 | 6.5 | 223 | 46.1 |
| G28881A,<br>G28882A,<br>G28883C | N | R203K,<br>G204R | 2 | 1.0 | 73 | 15.1 |
| A29827T | 3' UTR | - | 0 | 0.0 | 44 | 9.1 |
| G29830T | 3' UTR | - | 0 | 0.0 | 61 | 12.6 |

This difference in the mutational profile of isolates from Gujarat with that of the RoI is clearly seen in the diversity plots for non-synonymous mutations in isolates in Figure 4 (a) Gujarat, (b) Telangana and (c) India. It may be noted from plots 4(a) and 4(c) that the frequency of mutations in Gujarat isolates is very different from that of isolates from the whole of India. In contrast, the diversity plots of Telangana (Fig 4(b)) and India (Fig 4(c)) are quite similar, probably due to subclade I/A3i. The death rates for Telangana and India are also comparable, while that of Gujarat is strikingly different. This analysis suggests that the characteristic mutations of subclade I/A3i may not be deleterious.

**Fig. 4.**
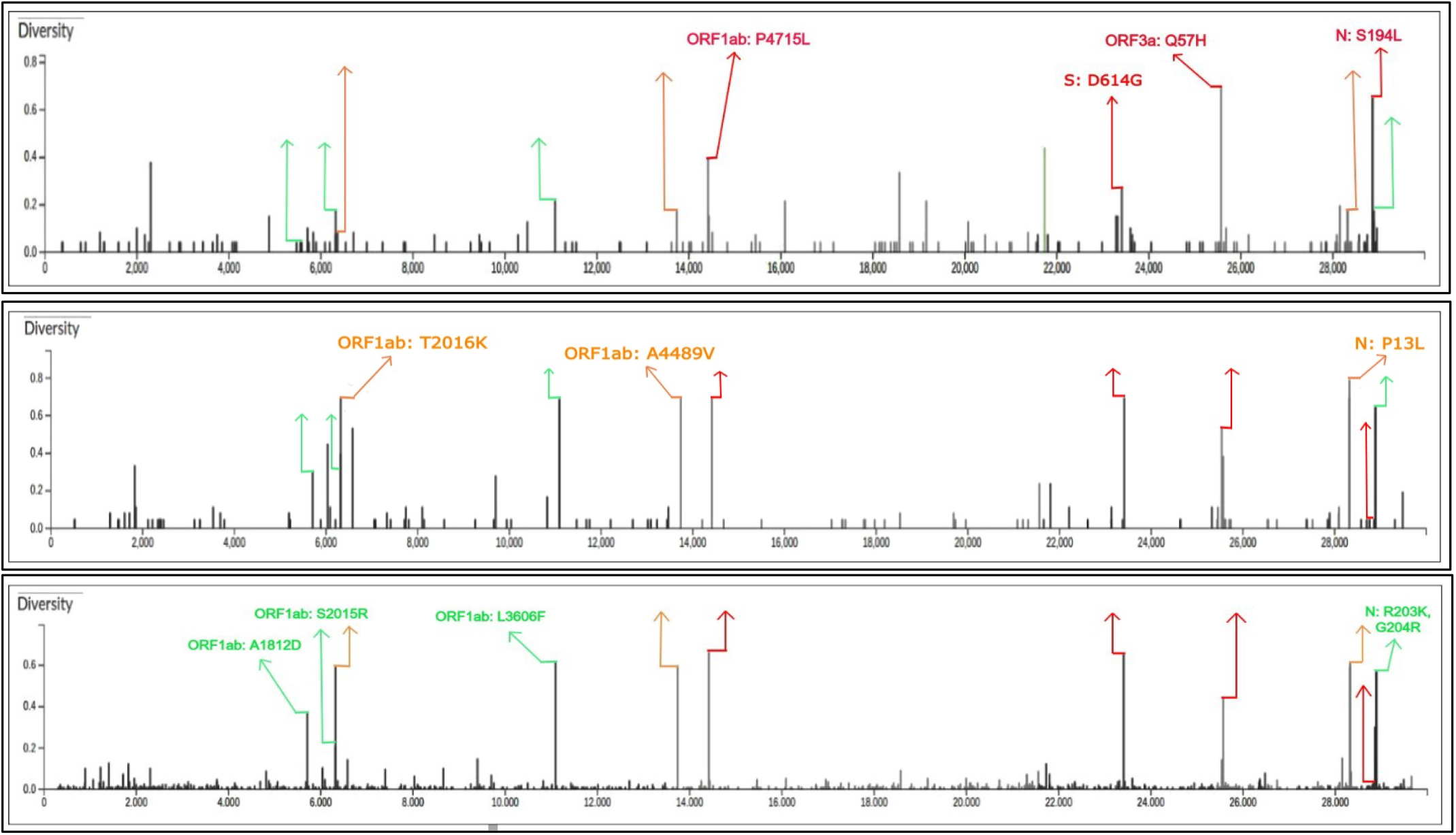
Diversity plots for non-synonymous mutations in isolates from (a) Gujarat, (b) Telangana, and (c) India clearly exhibit different sets of mutations.

### Identification of novel subclade I/GJ-20A and unique mutations in Maharashtra

Analysis of Gujarat isolates revealed a novel subclade, I/GJ-20A of clade 20A (**Supplementary Figure 2c**), defined by the shared mutations C18877T, G25563T (ORF3a: Q57H), and C26735T (listed in Table 2) in accordance with an earlier study (Joshi *et al*. 2021). These mutations are under-represented in RoI and their analysis may be helpful in explaining high death rates in Gujarat compared to the other Indian states. The Q57H mutation caused by G25563T is present on the outer surface of ORF3a protein and has been implicated in inflammation, antiviral responses, and apoptosis control. Mutations in ORF3a are of significance as the ORF3a protein is involved in regulating immunological responses in the host, including “cytokine storm”. The mutation results in significant conformational changes to the protein and forms a more stable quaternary structure thereby increasing the viral particle release (Wang *et al*. 2020; Oulas *et al*. 2021). It is indicated to increase the transmission.

According to I-Mutant2.0 and MUpro analysis, Q57H mutation decreases the stability of the protein. Other mutations that are part of this subclade are C2836T, C22444T, and C28854T (N: S194L). The S194L mutation due to C28854T resides in the central region of the nucleocapsid protein that is essential for oligomerization (Zhao *et al*. 2005; Yu *et al*. 2006) and is expected to alter the protein structure (Wu *et al*. 2021). It is predicted to increase protein stability by both I-Mutant2.0 and MUpro (Table 1).

A similar analysis of Maharashtra isolates when compared to isolates from RoI showed noticeable differences at the genetic level (Table 3). Maharashtra with second highest death rate (4.19%) after Gujarat also recorded highest number of infections. Two co-occurring mutations in the 3’ UTR, A29827T (**Supplementary Figure 2d**) and G29830T, exhibited a prevalence of 55% and 73.75% respectively in the isolates from Maharashtra but their presence was observed in only 2 isolates from RoI. Another set of co-occurring mutations, C313T and C5700A (ORF1ab: A1812D), was observed in Maharashtra with a high frequency of 25% and 26.25% respectively but with <3% in isolates from RoI. These two distinct sets of mutations form two distinct clusters of samples from Maharashtra in the PCA plot. Among these only C5700A (A1812D) is a non-synonymous mutation in the Nsp3 protein of ORF1ab and its functional significance is discussed above.

**Table 3.** Mutations observed in SARS-COV-2 isolates from Maharashtra (80 samples) with high frequency compared to the Rest of India (605 samples) are listed.

| Nucleotide Mutation | Protein | Amino Acid Mutation | Count in Maharashtra | Frequency in Maharashtra (%) | Count in RoI | Frequency in RoI (%) |
| --- | --- | --- | --- | --- | --- | --- |
| C313T | ORF1ab (nsp1) | - | 20 | 25.0 | 19 | 3.1 |
| C5700A | ORF1ab (nsp3) | A1812D | 21 | 26.2 | 12 | 2.0 |
| A29827T | 3' UTR | - | 44 | 55.0 | 0 | 0.0 |
| G29830T | 3' UTR | - | 59 | 73.8 | 2 | 0.3 |

## Conclusion

In this study, demographic analysis of mutations in Indian SARS-CoV-2 isolates was conducted to understand the viral spread in the country during early phase of the pandemic (27^th^ Jan 2020 – 27^th^ May 2020) and assess the effectiveness of contact tracing, quarantine and lockdown in controlling its spread. Genetic analysis revealed that though lockdown helped in controlling the spread of the virus, region-specific set of shared mutations observed indicate local transmission within the states. This provides probable explanation of observed variation in the number of infected cases and death rates across states. Region specific sequencing efforts can help in understanding the mutational spectra in local hotspots like Gujarat and Maharashtra and aid in the implementation of state-wise lockdown policies. Due to limited sequencing data, numerous variants are likely to have been missed. A vast country like India needs to improve its sequencing efforts to understand transmission dynamics of the virus, capture novel variants and assess their clinical impact to follow appropriate containment measures; this would prepare us for the third and fourth waves of COVID-19.

## Data Availability statement

The data that support the findings of this study are openly available in GISAID at https://www.gisaid.org/.

## Supplementary Figures

**Supplementary Fig. 1**. State-wise distribution of the clades is shown. Clade 20A is predominant in Gujarat while the root clade 19A is observed majorly in states Telangana, Delhi, and Maharashtra.

**Supplementary Fig. 2**. Sequences carrying the mutations a) C5700A b) C23929T c) 18877T d) G29830T are depicted in yellow colour.

**Supplementary Fig. 3 (a)** State-wise distribution of subclade I/A3i isolates in India. **(b)** The global distribution of the isolates with C23929T mutation (representative of I/A3i subclade) is shown (zoomed into Asia).

## References

Alam A. S. M. R. U., Islam O. K., Hasan M. S., Islam M. R., Mahmud S., Al-Emran H. M. et al. 2021 Dominant Clade-featured SARS-CoV-2 Co-occurring Mutations Reveals Plausible Epistasis: An in silico based Hypothetical Model. medRxiv.

Ayub M. I. 2020 Reporting Two SARS-CoV-2 Strains Based on A Unique Trinucleotide-Bloc Mutation and Their Potential Pathogenic Difference. Preprints.

Bai Y., Jiang D., Lon J. R., Chen X., Hu M., Lin S. et al. 2020 Comprehensive evolution and molecular characteristics of a large number of SARS-CoV-2 genomes reveal its epidemic trends. J. Infect. Dis. 100, 164–173.

Banu S., Jolly B., Mukherjee P., Singh P., Khan S., Zaveri L. et al. 2020 A Distinct Phylogenetic Cluster of Indian Severe Acute Respiratory Syndrome Coronavirus 2 Isolates. Open Forum Infect. Dis. 7, ofaa434.

Capriotti E., Fariselli P. and Casadio R. 2005 I-Mutant2.0: Predicting stability changes upon mutation from the protein sequence or structure. Nucleic Acids Res. 33(Web Server issue), W306–310.

Chang C.-K., Hsu Y.-L., Chang Y.-H., Chao F.-A., Wu M.-C., Huang Y.-S. et al. 2009 Multiple nucleic acid binding sites and intrinsic disorder of severe acute respiratory syndrome coronavirus nucleocapsid protein: Implications for ribonucleocapsid protein packaging. J. Virol. 83, 2255–2264.

Chang C., Hou M.-H., Chang C.-F., Hsiao C.-D. and Huang T. 2014 The SARS coronavirus nucleocapsid protein—Forms and functions. Antivir. Res. 103, 39–50.

Chaudhari A., Chaudhari M., Mahera S., Saiyed Z., Nathani N. M., Shukla S. et al. 2021 In-Silico analysis reveals lower transcription efficiency of C241T variant of SARS-CoV-2 with host replication factors MADP1 and hnRNP-1. Inform. Med. Unlocked 25, 100670.

Cheng J., Randall A. and Baldi P. 2006 Prediction of protein stability changes for single-site mutations using support vector machines. Proteins 62, 1125–1132.

Daniloski Z., Jordan T. X., Ilmain J. K., Guo X., Bhabha G., tenOever B. R. et al. 2021 The Spike D614G mutation increases SARS-CoV-2 infection of multiple human cell types. Elife 10, e65365.

Gao Y., Yan L., Huang Y., Liu F., Zhao Y., Cao L. et al. 2020 Structure of the RNA-dependent RNA polymerase from COVID-19 virus. Science 368, 779–782.

Gupta A., Sabarinathan R., Bala P., Donipadi V., Vashisht D., Katika M. R. et al. 2021 A comprehensive profile of genomic variations in the SARS-CoV-2 isolates from the state of Telangana, India. J. Gen. Virol. 102.

Hadfield J., Megill C., Bell S. M., Huddleston J., Potter B., Callender C. et al. 2018 Nextstrain: Real-time tracking of pathogen evolution. Bioinformatics 34, 4121–4123.

Hsin W.-C., Chang C.-H., Chang C.-Y., Peng W.-H., Chien C.-L., Chang M.-F. et al. 2018 Nucleocapsid protein-dependent assembly of the RNA packaging signal of Middle East respiratory syndrome coronavirus. J. Biomed. Sci. 25, 47.

Joshi M., Puvar A., Kumar D., Ansari A., Pandya M., Raval J. et al. 2021 Genomic Variations in SARS-CoV-2 Genomes From Gujarat: Underlying Role of Variants in Disease Epidemiology. Front. Genet. 12, 292.

Korber B., Fischer W. M., Gnanakaran S., Yoon H., Theiler J., Abfalterer W. et al. 2020 Tracking Changes in SARS-CoV-2 Spike: Evidence that D614G Increases Infectivity of the COVID-19 Virus. Cell 182, 812–827.e19.

Lei J., Kusov Y. and Hilgenfeld R. 2018 Nsp3 of coronaviruses: Structures and functions of a large multi-domain protein. Antivir. Res. 149, 58–74.

Maitra A., Sarkar M. C., Raheja H., Biswas N. K., Chakraborti S., Singh A. K. et al. 2020 Mutations in SARS-CoV-2 viral RNA identified in Eastern India: Possible implications for the ongoing outbreak in India and impact on viral structure and host susceptibility. J. Biosci. 45, 76.

Masters P. S. 2019 Coronavirus genomic RNA packaging. Virol. J. 537, 198–207.

Oulas A., Zanti M., Tomazou M., Zachariou M., Minadakis G., Bourdakou M. M. et al. 2021 Generalized linear models provide a measure of virulence for specific mutations in SARS-CoV-2 strains. PLoS One 16, e0238665.

Paul D., Jani K., Kumar J., Chauhan R., Seshadri V., Lal G. et al. 2020 Phylogenomic analysis of SARS-CoV-2 genomes from western India reveals unique linked mutation. bioRxiv.

Plante J. A., Liu Y., Liu J., Xia H., Johnson B. A., Lokugamage K. G. et al. 2021 Spike mutation D614G alters SARS-CoV-2 fitness. Nature 592, 116–121.

Shu Y. and McCauley J. 2017 GISAID: Global initiative on sharing all influenza data - from vision to reality. Euro Surveill. 22, 30494.

Tylor S., Andonov A., Cutts T., Cao J., Grudesky E., Van Domselaar G. et al. 2009 The SR-rich motif in SARS-CoV nucleocapsid protein is important for virus replication. Can. J. Microbiol. 55, 254–260.

Wang R., Chen J., Gao K., Hozumi Y., Yin C. and Wei G. 2020 Characterizing SARS-CoV-2 mutations in the United States. Res. Sq., rs.3.rs-49671.

Wu S., Tian C., Liu P., Guo D., Zheng W., Huang X. et al. 2021 Effects of SARS-CoV-2 mutations on protein structures and intraviral protein-protein interactions. J. Med. Virol. 93, 2132–2140.

Yu I.-M., Oldham M. L., Zhang J. and Chen J. 2006 Crystal structure of the severe acute respiratory syndrome (SARS) coronavirus nucleocapsid protein dimerization domain reveals evolutionary linkage between corona-and arteriviridae. J. Biol. Chem. 281, 17134–17139.

Zhang L., Jackson C. B., Mou H., Ojha A., Peng, H. Quinlan B. D. et al. 2020 SARS-CoV-2 spike-protein D614G mutation increases virion spike density and infectivity. Nat. Commun. 11, 6013.

Zhao P., Cao J., Zhao L.-J., Qin Z.-L., Ke J.-S., Pan W. et al. 2005 Immune responses against SARS-coronavirus nucleocapsid protein induced by DNA vaccine. Virol. J. 331, 128–135.

