## Supplementary material for "Demographic Analysis of Mutations in Indian SARS-CoV-2 Isolates": Supplrementary Figure

**Supplementary Figures**

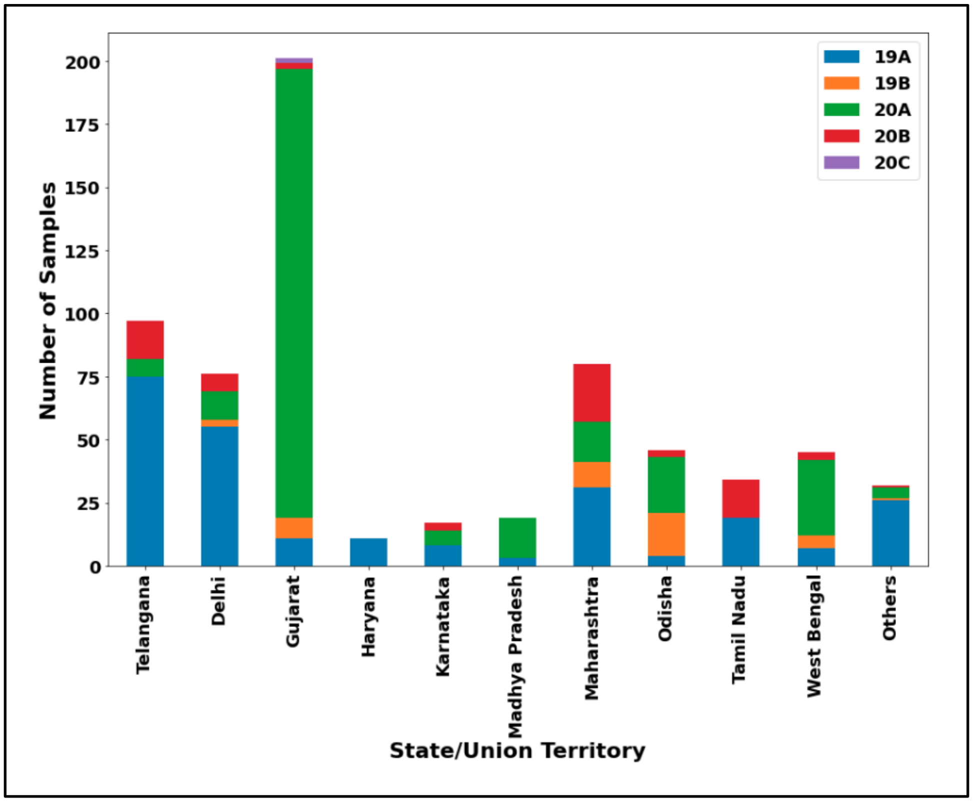


**Supplementary Fig. 1.** State-wise distribution of clades. Clade 20A is predominant in Gujarat while the root clade 19A is observed majorly in states Telangana, Delhi, and Maharashtra.

(a)

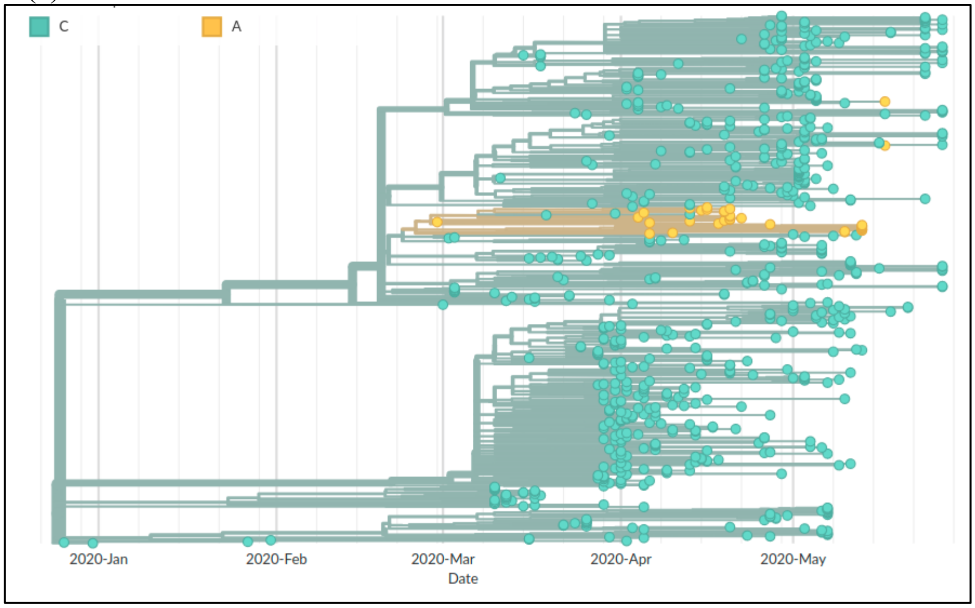

(b)

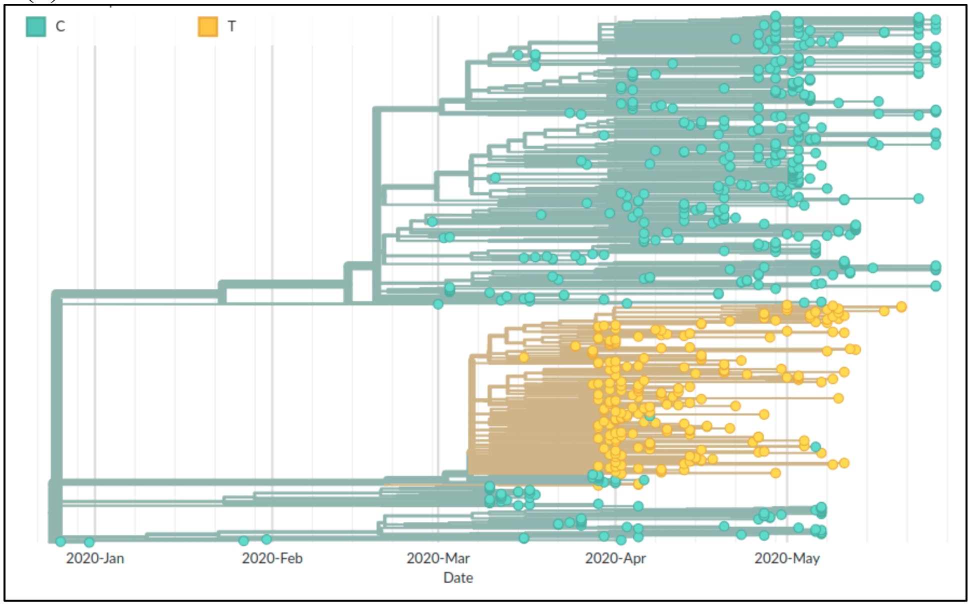
C
T
2020-Jan
2020-Feb
2020-Mar
Date
2020-Apr
2020-May

(c)

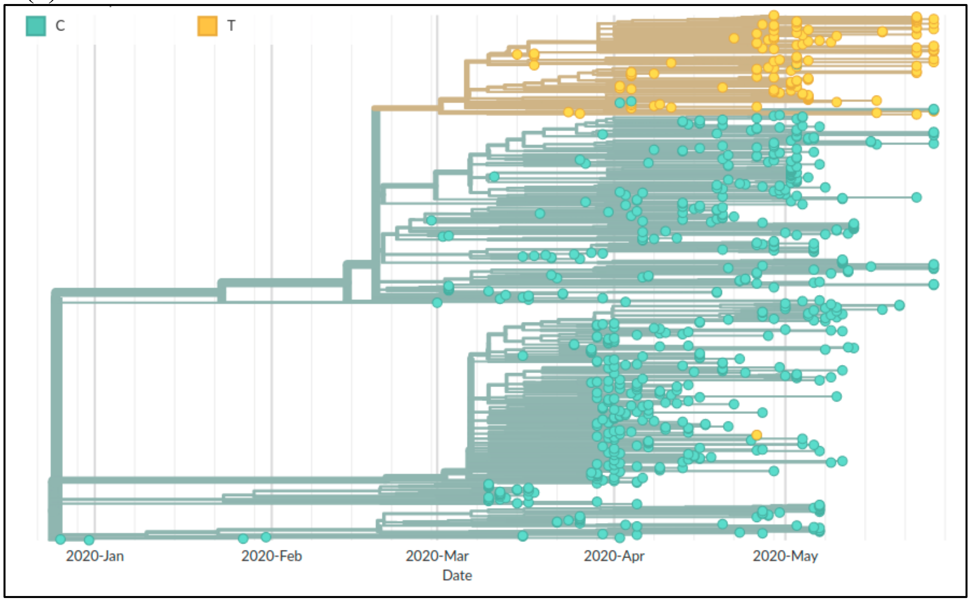
C
T
2020-Jan
2020-Feb
2020-Mar
Date
2020-Apr
2020-May

(d)

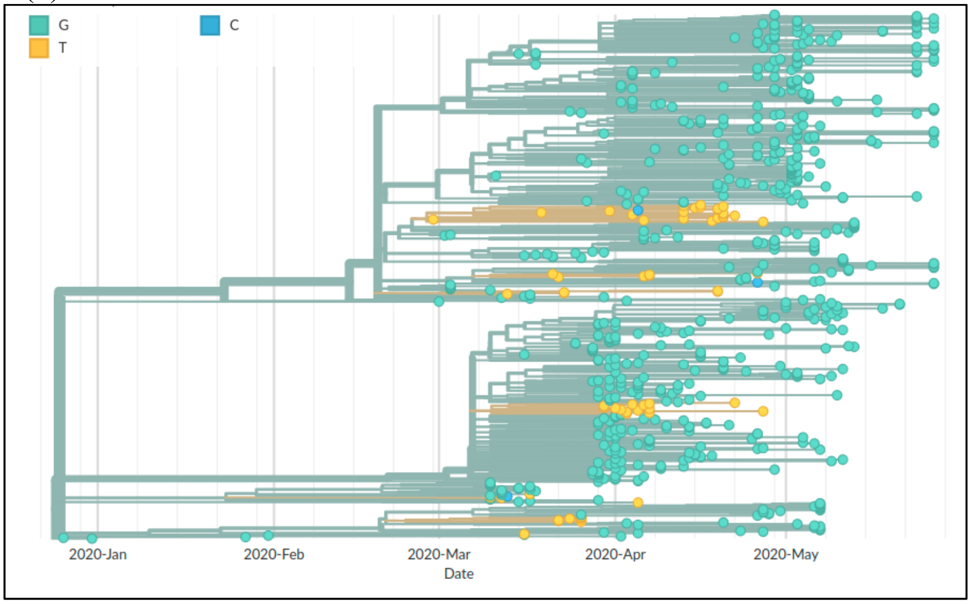


**Supplementary Fig. 2.** The sequences carrying the mutations a) C5700A b) C23929T c) 18877T d) A29827T are depicted in yellow colour.

(a)

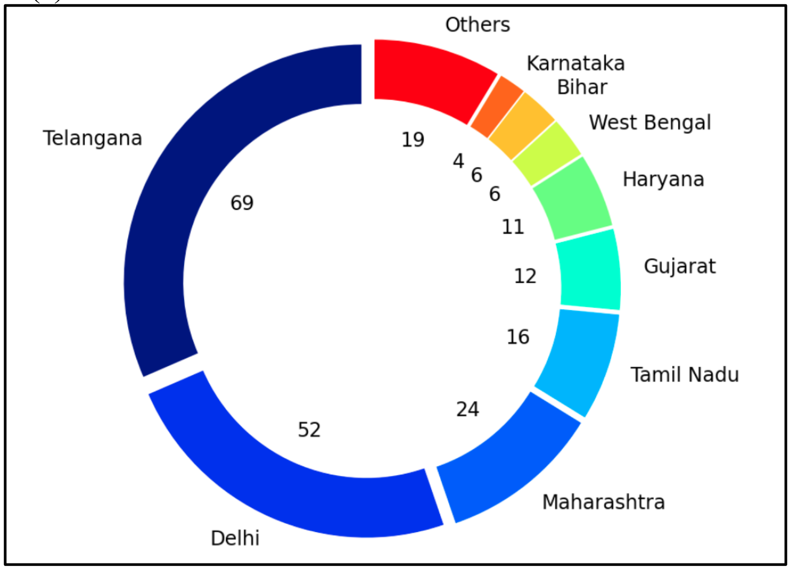

(b)

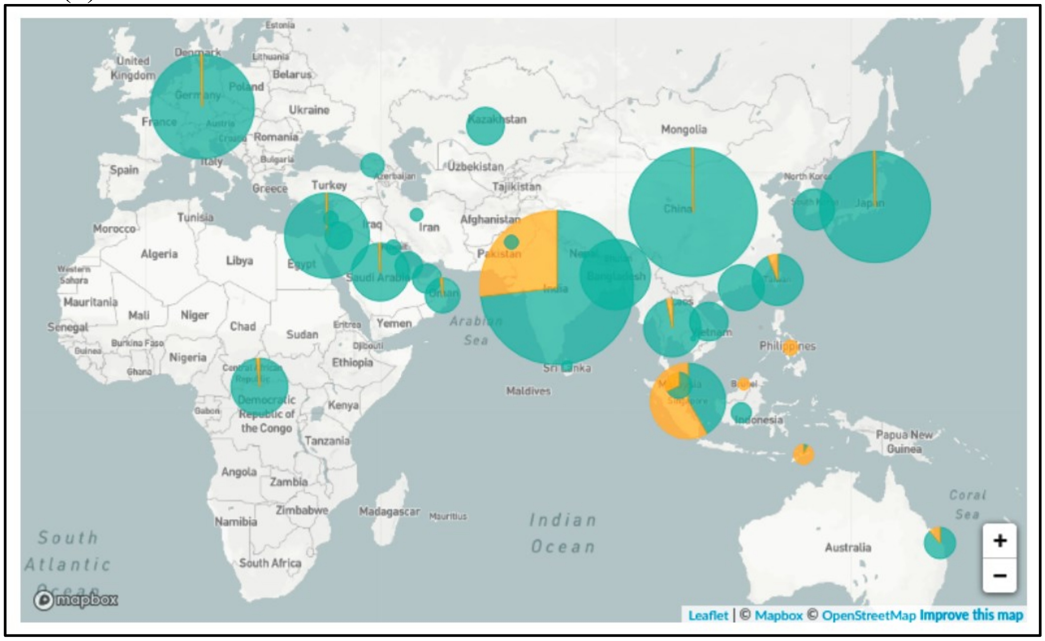


**Supplementary Fig. 3 (a)** State wise distribution of the subclade I/A3i isolates in India. **(b)** The global distribution (zoomed in to Asia) of the mutation C23929T part of I/A3i is shown.
